# Ammonia-based enrichment and long-term propagation of zone I hepatocyte-like cells

**DOI:** 10.1101/2020.03.20.999680

**Authors:** Ruri Tsuneishi, Noriaki Saku, Shoko Miyata, Palaksha Kanive Javaregowda, Kenta Ite, Masashi Toyoda, Tohru Kimura, Masahiko Kuroda, Atsuko Nakazawa, Mureo Kasahara, Hidenori Nonaka, Akihide Kamiya, Tohru Kiyono, Junji Yamauchi, Akihiro Umezawa

## Abstract

Zone I and zone III hepatocytes metabolize ammonia through urea cycle and drug by cytochrome P450, respectively. Ammonia has a cytotoxic effect, and can therefore be used as a selection agent for enrichment of hepatocytes. Besides, isolated hepatocytes from livers can be propagated ex vivo under appropriate condition. However, it has not been investigated so far whether ammonia-treated hepatocyte-like cells are able to proliferate in vitro. In this study, we employed the ammonia selection strategy to purify hepatocyte-like cells that were differentiated from human pluripotent stem cells (PSCs) that are embryonic stem cells (ESCs) and induced pluripotent stem cells. Hepatocyte-like cells after exposure to ammonia highly expressed the CPS1 gene that metabolizes ammonia to carbamoyl phosphate. The resistance to cytotoxicity or cell death by ammonia is probably attributed to the metabolic activity of ammonia in the cells. In addition to the ammonia metabolism-related genes, ammonia-selected PSC-derived hepatocytes increased expression of the CYP3A4 gene, one of the cytochrome P450 genes, that is mainly expressed in zone III hepatocytes. Ammonia-selected hepatocyte-like cells derived from both ESCs and iPSCs can be propagated in vitro up to 30 population doublings for more than 190 days without affecting expression of the liver-associated genes, implying that the ammonia-selected cells have immortality or equivalent life span on the appropriate feeder cells like ESCs and iPSCs. The long-term cultivation of ammonia-selected hepatocyte-like cells resulted in the increased expression of hepatocyte-associated genes such as the CPS1 and CYP3A4 genes. The ammonia selection method to enrich a hepatocyte population was also applicable to immortalized cells from the liver. Ammonia treatment in combination with in vitro propagation will be used to obtain large amounts of hepatocytes or hepatocyte-like cells for pharmacology, toxicology and regenerative medicine.

## Introduction

Hepatocytes have been used as a substitute of animal experiments for preclinical safety tests to investigate hepatotoxicity of low-molecular drugs [1]. Primary culture of hepatocytes dissociated from a liver is a golden standard in pharmaceutical in vitro studies for clinical prediction. The drawbacks of the use of hepatocytes in drug screening includes limited supply of the same lot and large variations between lots due to genetic and environmental backgrounds. To solve the lot variation issue, HepG2 is also used to examine hepatotoxicity because of its clonal nature. HepaRG, another hepatocyte-like clone, was established from hepatoblastoma, and has an advantage in high induction of cytochrome P450 genes [2]. In addition to isolated hepatocytes, immortalized hepatocytes, and hepatocarcinoma cells, hepatocytes or hepatocyte-like cells can be differentiated from human pluripotent stem cells (PSCs) such as embryonic stem cells (ESCs) and induced pluripotent stem cells (iPSCs). Human PSCs impact numerous medical fields including clinical therapy development, drug discovery, research on inherited diseases and studies on reprogramming of differentiated cells [3-6]. For example, human PSC-derived hepatocytes serve as an in vitro tool for understanding drug metabolism and toxicology [7,8]. Human PSC-derived hepatocytes or hepatocyte-like cells can be obtained from the same origin repeatedly due to immortality of ESCs [9-11]. Parenchymal hepatocytes in liver lobules have different characteristics and function in lobular zones: Zone I hepatocytes around portal area, zone III hepatocytes around central vein, and zone II hepatocytes that intermediarily locate. Zone I hepatocytes have metabolic activity of ammonia and glycogenesis, while zone III hepatocytes have drug metabolic activity and glycolysis. The zoning is considered to depend on concentrations of oxygen and hormones. Hepatocyte-like cells differentiated from human PSCs are enriched with ammonia because these cells have a high metabolic activity of ammonia that is toxic to mammalian cells, especially neural cells [12]. Hepatocytes can be maintained in vitro without any introduction of any genes [13,14]. In this study, we propagated hepatocytes or hepatocyte-like cells derived from human PSC, i.e. ESCs and iPSCs, and enriched zone I hepatocytes with ammonia [15,16]. Ammonia-selected cells proliferated in vitro at least up to 30 population doublings for more than 190 days. This study introduced an alternative strategy to obtain enough number of human hepatocytes with a combination of ammonia selection and subsequent in vitro propagation.

## Results

### Selection of ESC-derived hepatocyte-like cells with ammonia

To obtain zone I hepatocytes, SEES2 ESCs were differentiated into hepatocyte-like cells (Figure 1A-D). After differentiation, ESC-derived hepatocyte-like cells were exposed to ammonia for 2 days. The heterogenous population of the differentiated cells was killed with ammonia at 70 to 80%. The ammonia-selected cells started to proliferate as colonies and exhibited hepatocyte-like morphology.

**Figure 1.**
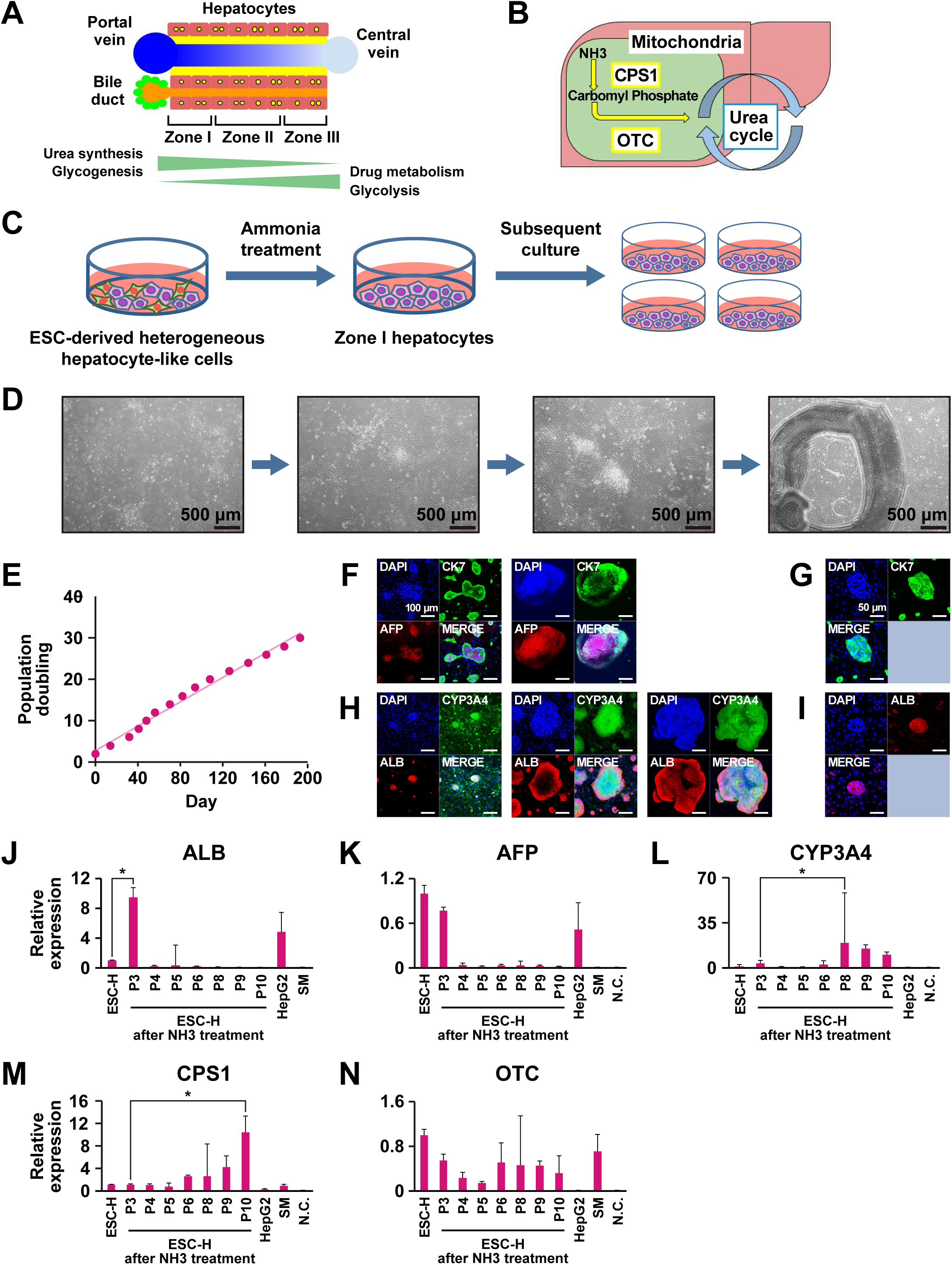
Generation of hepatocyte-like cells from SEES2 ESCs after exposure to ammonia. (A-B) Hepatocytes with ammonia metabolism in liver. (C) Scheme for characterization of ammonia-selected hepatocyte-like cells and ammonia metabolism. (D) Phase-contrast photomicroscopy of the hepatocyte-like cells during ammonia selection. Bar:500μm. (E) Growth curve of the ammonia-selected cells. (F) Immunocytochemistry of the ammonia-selected cells by using the antibodies to CK7 and AFP. (G) Immunocytochemistry of the ammonia-selected cells by using the antibodies to CK7. (H) Immunocytochemistry of the ammonia-selected cells by using the antibodies to CYP3A4 and ALB. (I) Immunocytochemistry of the ammonia-selected cells by using the antibodies to ALB. (J-N) qRT-PCR analysis of the genes for ALB (J), AFP (K), CYP3A4 (L), CPS1 (M) and OTC (N). From left to right: ESC-derived hepatocyte-like cells without ammonia selection,ammonia-selected ESC-derived hepatocyte-like cells at passage 3-10. HepG2 cells, negative control (H_2_O), iPSC-derived immortalized hepatocyte-like cells (HepaSM).

### Long-term cultivation of ESC-derived hepatocyte-like cells

After exposure to ammonia, the cells escaped from cell death retained replicative capacity on MEF feeder. The ammonia-selected cells continued to proliferate in vitro at least up to 30 population doublings for more than 190 days (Figure 1E). Immunocytochemistry revealed that the ammonia-selected cells were positive for AFP, ALB, CYP3A4, and CK7 (Figure 1F-I). Immnoreactivity of the ammonia-selected cells at the central and peripheral portion of the colonies are relatively strong for AFP and CK7, respectively (Figure 1F). The cells at the peripheral portion of colonies stained positive for ALB, and almost all cells in the colonies stained positive for CYP3A4 (Figure 1H). In certain colonies, all cells in colonies are positive for CK7 and ALB (Figure 1G, I). The results imply that the ammonia-selected cells retained hepatic and ductal characteristics even at more than 30 PDs. The ammonia-selected hepatocyte-like cells were then applied to qRT-PCR analysis to investigate expression levels of hepatocyte-associated genes (Figure 1J-N). The ammonia-selected cells just after selection at 3 passages expressed the genes for ALB, AFP, CYP3A4, CPS1, and OTC. The expression levels of the genes for ALB and AFP decreased, but those of the genes for CYP3A4 and CPS1 dramatically increased during the serial passaging. The expression levels of the OTC gene remained unchanged upon passaging.

### Feeder cells for in vitro proliferation of ESC-derived hepatocyte-like cells

Proliferation of the ESC-derived ammonia-selected cells was dependent on MEF feeder cells. The ammonia-selected cells started to detach from dishes without MEF feeder in 2 weeks. We therefore compared the ammonia-selected cells on the MEF feeder and feeder-free condition (Figure 2A). The ammonia-selected cells steadily proliferated on the MEF feeder condition (Figure 2B). Expression levels of the genes for ALB, AFP, CPS1, OTC and CK7 were higher on the MEF condition than on the feeder-free condition (Figure 2C-G).

**Figure 2.**
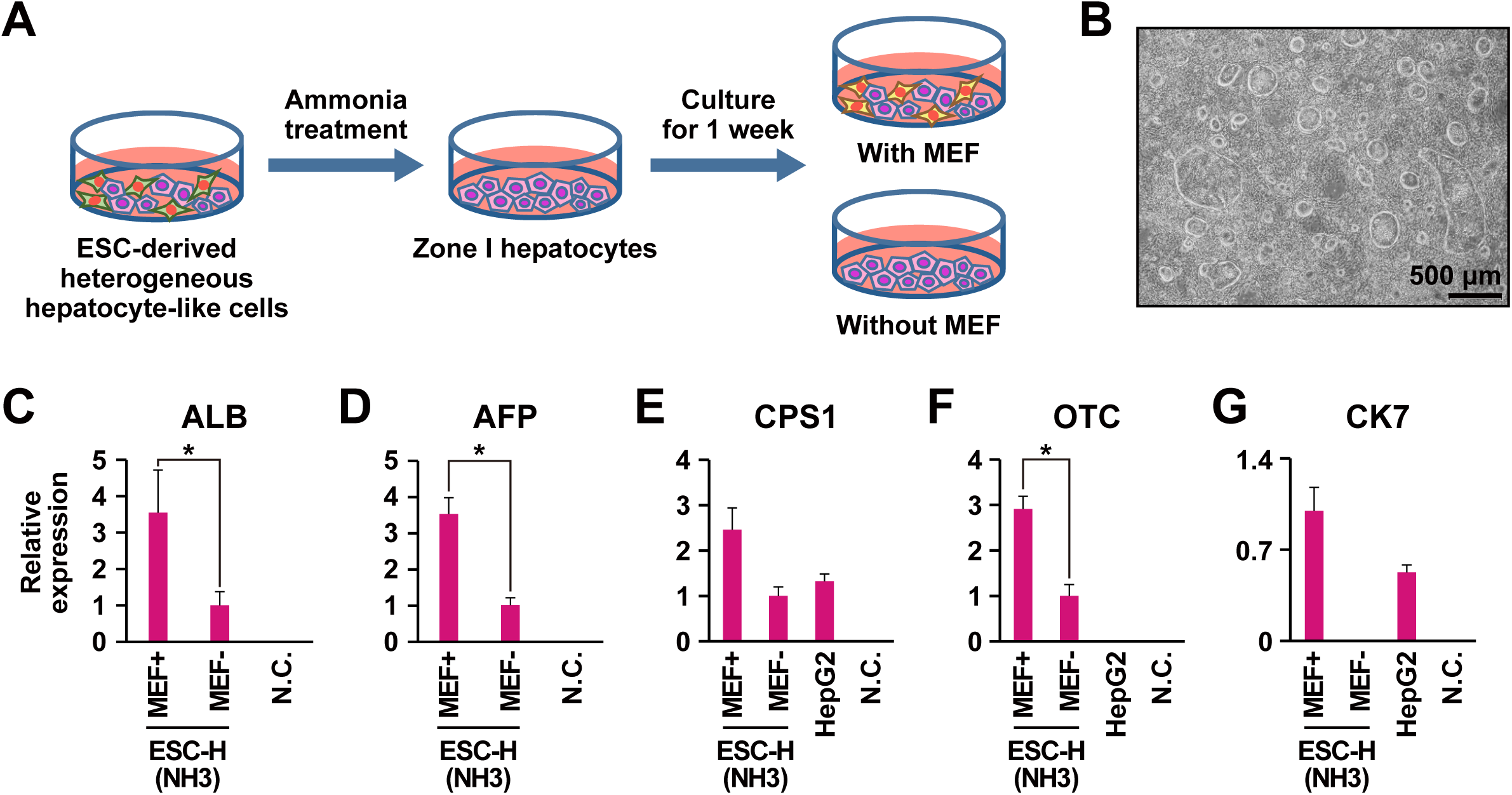
Requirement of feeder cells to support in vitro proliferation of the ammonia-selected hepatocyte-like cells. (A) Scheme for feeder cell dependency of ammonia-selected hepatocyte-like cells. (B) Phase contrast photomicrograph of the ammonia-selected hepatocyte-like cells on MEFs. (C-G) qRT-PCR analysis of the genes for ALB (C), AFP (D), CPS1 (E), OTC (F) and CK7 (G).

### Spheroid formation of the ESC-derived hepatocyte-like cells for maturation

We examined the ammonia-selected cells to investigate hepatocytic maturation through spheroid formation. We employed the 3D culture method to form spheroid (Figure 3A). Spheroids became visible on Day 3, formed on Day 6, and remained unchanged on Day 9 (Figure 3B). The genes for AFP, OTC and CYP3A4 were clearly up-regulated in the cells through the spheroid formation (Figure 3D-H).

**Figure 3.**
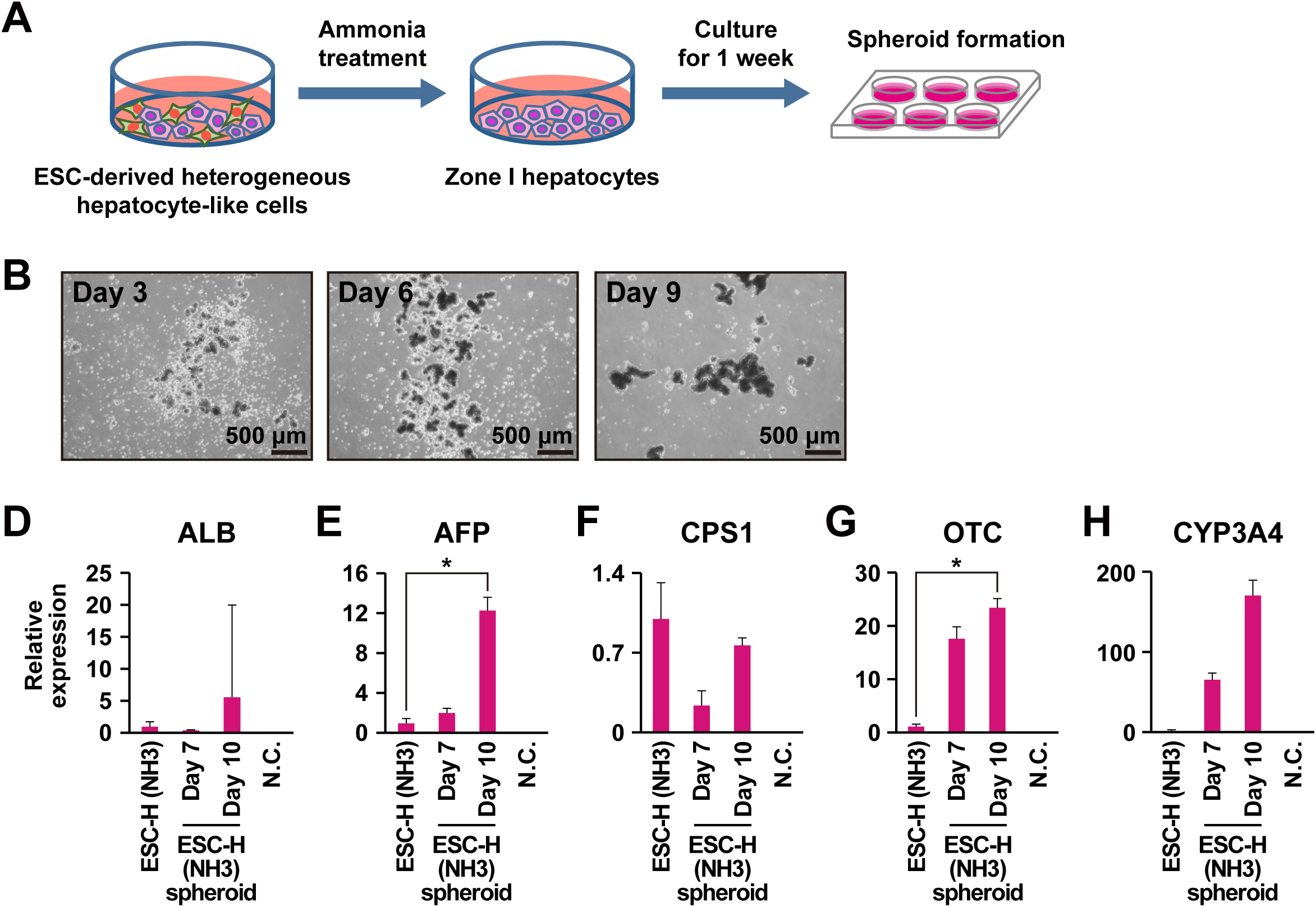
Up-regulation of the genes for ammonia metabolism via three-dimensional cultivation in the ammonia-selected hepatocyte-like cells. (A) Protocol for sphere formation of ammonia-selected hepatocyte-like cells. (B) Phase-contrast photomicrograph of the ammonia-selected hepatocyte-like cells at 3, 6, and 9 days after the sphere formation. (D-H) qRT-PCR analysis of the genes for ALB (D), AFP (E), CPS1 (F), OTC (G) and CYP3A4 (H). From left to right: Ammonia-selected hepatocyte-like cells at passage 12 without sphere formation, ammonia-selected hepatocyte-like cells at passage 12 with sphere formation (Day7,Day10), HepG2 cells, negative control (H_2_O).

### Selection of iPSC-derived hepatocyte-like cells with ammonia

In addition to ESC differentiation, iPSCs were examined to have a potency to differentiate into hepatocytes. We used iPSC-O i.e. iPSCs from a patient with drug-induced hepatic injury (DILI-iPSCs), for hepatic differentiation. iPSC-O cells were exposed to ammonia for 2 days after they were differentiated into heterogenous population of differentiated cells (Figure 4A). The differentiated cells including hepatocyte-like cells were killed with ammonia in three independent experiments (Figure 4B-D). The cell-killing effect of ammonia depended on ammonia concentration. After the exposure to ammonia, the selected cells exhibited hepatocyte-like morphologies such as epithelial-monolayered flatter cells, a large cytoplasmic-to-nuclear ratio, numerous and prominent nucleoli, and occasional binucleated cells. These hepatocyte-like cells were propagated in vitro and examined for expression of the hepatocyte-associated genes (Figure 4E-I). The iPSC-O differentiated cells after the ammonia selection increased the expression of the genes for ALB, AFP, and OTC at 2.1-, 20.1-, and 3.0-fold, respectively.

**Figure 4.**
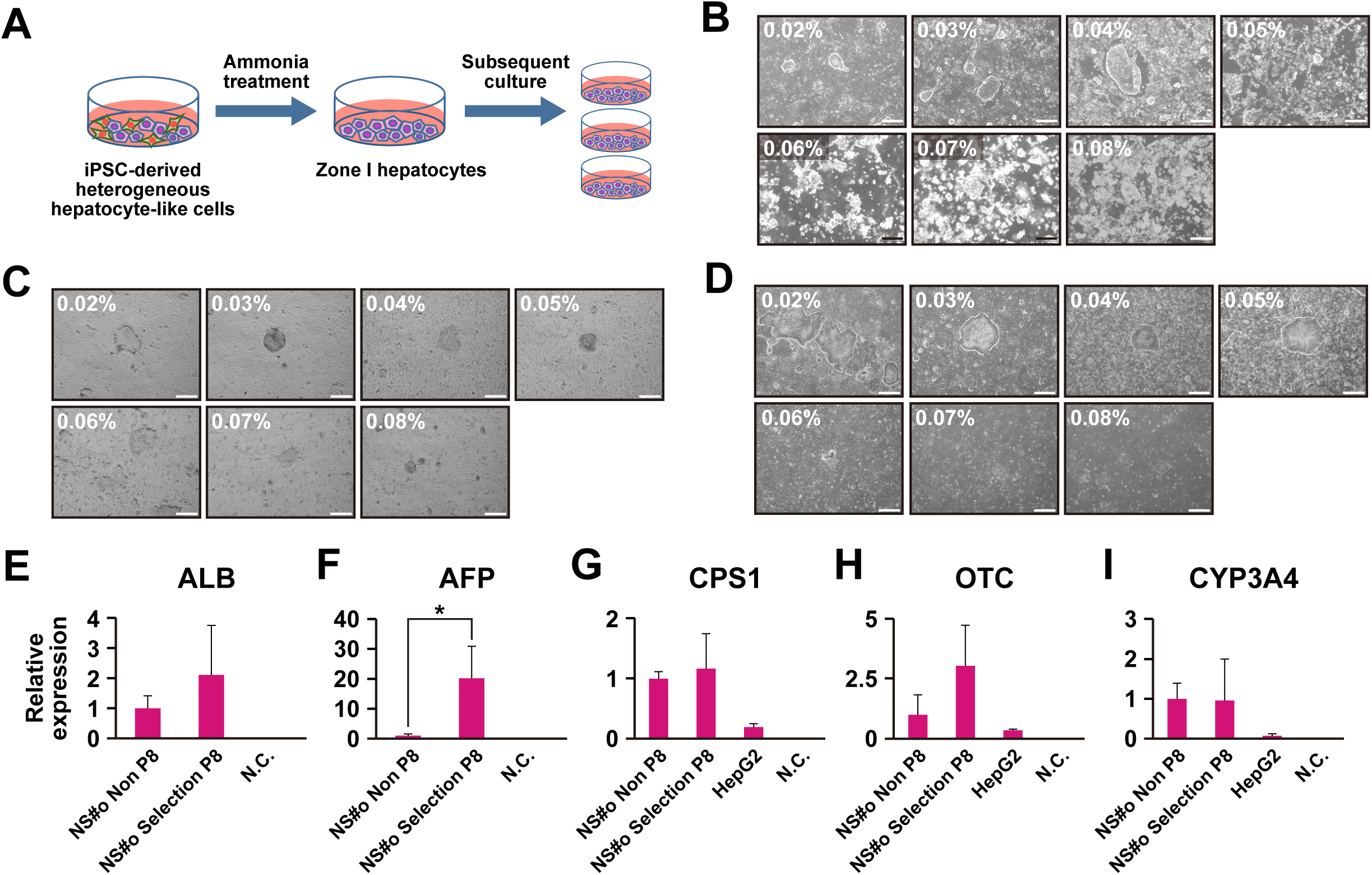
Ammonia selection of iPSC-derived hepatocyte-like cells. (A) Scheme for ammonia selection of iPSC-derived hepatocyte-like cells. (B-D) Phase contrast photomicrograph of the hepatocyte-like cells generated from DILI-derived iPSC-O cells exposure to the indicated concentration of ammonia for 0 day (B), and with exposure to the indicated concentration of ammonia for 1 day (C) or 2 days (D). (E-I) qRT-PCR analysis of the genes for ALB (E), AFP (F), CSP1 (G), OTC (H) and CYP3A4 (I). From left to right: Hepatocyte-like cells generated from DILI-derived iPSC-O cells without ammonia selection, ammonia-selected hepatocyte-like cells generated from DILI-derived iPSC-O cells, HepG2 cells, negative control (H_2_O).

## Discussion

### Ammonia-based enrichment of zone I hepatocytes

Ammonia has a cell-killing effect; however, the mechanism remains unclarified. Ammonium ions compete with potassium ions for inward transport, over the cytoplasmic membrane, via potassium transport proteins like the Na+/K +ATPase and the Na+K+2Cl-cotransporter. This competition of ammonia with potassium ions lead to predictable intracellular and/or extracellular pH change and subsequent cell death. Neural toxicity of ammonia is associated with intracellular pH change through inhibition of potassium channel and depolarization of GABA neuron [17,18]. Hepatocytes predictably have high metabolic activity of ammonia and can escape from cytotoxic ammonia. The escape of PSC-derived differentiated cells from ammonia is therefore explained by endogenous ammonia metabolic activity, i.e. expression of the urea cycle-related genes such as OTC and CPS1 [12]. To enrich cells with ammonia metabolic activity, selection with cell surface markers such as CD13 by flow cytometric analysis and magnetic cell sorting can be used. Introduction of cell type-specific expression of cytotoxic antibiotics-resistant genes such as the neomycin-resistant gene is also available for cell selection [20]. Compared with these sophisticated approach, successful enrichment with ammonia exposure used in this study is simple and straightforward. Indeed, the ammonia selection method was also applicable to not only PSC-derived cells but also HepaMN immortalized cell line. It is also noteworthy that ammonia selection may increase homogeneity or decrease heterogeneity among a lot from the viewpoint of ammonia metabolism activity.

### Altered and unaltered gene expression after ammonia selection and during propagation

It is of note that the levels of the hepatocyte-associated genes such as the CPS1 and CYP3A4 genes gradually increase during the long-term cultivation of ammonia-selected hepatocyte-like cells. Up-regulation of the CPS1 gene, one of the ammonia-metabolizing genes, after ammonia exposure and subsequent propagation is convincing because the increased expression might be correlated with ammonia metabolism activity. In contrast to urea cycle-related genes, up-regulation of the CYP3A4 gene, one of the cytochrome P450 genes, after ammonia selection and subsequent propagation is rather against our expectation because this gene is considered to be expressed in zone III hepatocytes [21]. Zone I and III hepatocytes are involved in ammonia metabolism and drug metabolism, respectively. This increase of the cytochrome P450 gene expression after enrichment of PSC-derived hepatocytes is probably explained by an increased number of the PSC-derived zone I hepatocytes with expression of the cytochrome P450 gene. Another possibility is that the ammonia-selected zone I hepatocytes start to express the cytochrome P450 genes during long-term cultivation in PSC-derived cells. Along with the increase of the cytochrome P450 genes, the ALBUMIN gene decreased during the cultivation. This reciprocal change of these genes can also be attributed to populational change of the cell types or change of gene expression in the cells. These changes in gene expression is not correlated with physiological condition or developmental process but could be an artifact of in vitro event. However, the cells with high ammonia metabolic activity and high CYP3A4 expression obtained in more than 200 days are obviously available for tests of pharmacokinetics/pharmacodynamics and toxicology, and can also be used for a model of regenerative therapy products.

### Effect of three-dimensional organoid formation on maturation

Maturation procedures of hepatocytes and hepatocytic progenitors include exposure of the cells with low molecules such as dexamethazon and demethylsulfoxide and cytokines such as Oncostatin M and hepatocyte growth factor. Cultivation in the B27 medium is also used for maturation. Contribution of the three-dimensional organoid formation to up-regulation of the genes for ALB, AFP and OTC was impressive because Oncostatin M and hepatocyte growth factor had a little effect on maturation. Spheroid is hard to use for toxicology tests because spheroid usually floats in the culture medium, but is preferred for regenerative therapy products if we consider the embolization of the spheroids carefully because of their high levels of the hepatocyte-associated markers.

### Immortality or limited replicative capacity

We are not able to conclude that the ammonia-selected hepatocyte-like cells derived from both ESCs and iPSCs have finite or infinite cell life span from this study. The life span of the ammonia-selected cells, up to 30 population doublings for more than 200 days, is enough to obtain sufficient amount of raw materials for drug development, pharmacokinetics, toxicology and regenerative therapy. Proof of immortality may require more than 100 population doublings, and this may be achieved by continuous cultivation for nearly 2 years, based on the cell growth curve (Figure 1E). Human hepatocytes are difficult to be propagated ex vivo due to lack of appropriate cultivation condition. To solve this issue, hepatocytes or hepatocyte-like cells have been obtained from livers, hepatoma/hepatocarcinoma, and ESCs/iPSCs [19,22-24]. Large-scale preparation of hepatocytes or hepatocyte-like cells with specific activities will be achieved with the three approaches: 1) in vitro propagation of cells with hepatocytic features, 2) maturation of propagated endodermal progenitors from iPSCs, 3) hepatic differentiation of propagated iPSCs.

### Possible regenerative therapy using ammonia-selected cells

Ammonia-selection and enrichment in combination with a strategy of generation and differentiation/maturation of human PSCs is one of the better strategies to take a robust system and a better cost-performance balance into a consideration. Hepatocytes have been implicated in cell-based therapy to patients with liver diseases or metabolic disorders who cannot receive liver transplantation. Likewise, human cells with a high ammonia metabolism activity generated from PSCs will be another source of cell therapy product. Patients with congenital metabolic disorders such as urea cycle disorder and citrullinemia suffer from hyperammonemia [19].

Hyperammonemia is caused by mutation or deletion of the ammonia metabolism-related genes. Ammonia-selected cells with a high ammonia metabolic activity may thus contribute the patients through an appropriate implantation strategy.

## Materials and Methods

### Ethical statement

Human cells in this study were performed in full compliance with the Ethical Guidelines for Medical and Health Research Involving Human Subjects (Ministry of Health, Labor, and Welfare, Japan; Ministry of Education, Culture, Sports, Science and Technology, Japan). The derivation and cultivation of ESC lines were performed in full compliance with “the Guidelines for Derivation and Distribution of Human Embryonic Stem Cells (Notification of the Ministry of Education, Culture, Sports, Science, and Technology in Japan (MEXT), No. 156 of August 21, 2009; Notification of MEXT, No. 86 of May 20, 2010) and “the Guidelines for Utilization of Human Embryonic Stem Cells (Notification of MEXT, No. 157 of August 21, 2009; Notification of MEXT, No. 87 of May 20, 2010)”. Animal experiments were performed according to protocols approved by the Institutional Animal Care and Use Committee of the National Research Institute for Child Health and Development.

### Culture of ESCs, iPSCs and hepatocytes

SEES” ESCs were routinely cultured onto a feeder layer of freshly plated gamma-irradiated mouse embryonic fibroblasts (MEFs), isolated from ICR embryos at 12.5 gestations and passages 2 times before irradiation (30 Gy), in the ESC culture media [15,16]. The ESC media consisted of Knockout™-Dulbecco’s modified Eagle’s medium (KO-DMEM) (Life Technologies, CA, USA; #10829-018) supplemented with 20% Knockout™-Serum Replacement (KO-SR; #10828-028), 2 mM Glutamax-I (#35050-079), 0.1 mM non-essential amino acids (NEAA; #11140-076), 50 U/ml penicillin-50 μg/ml streptomycin (Pen-Strep) (#15070-063), 0.055 mM β-mercaptoethanol (#21985-023) and recombinant human full-length bFGF (#PHG0261) at 10 ng/ml (all reagents from Life Technologies).

iPSC-O cells were generated from fibroblasts derived from a patient with DILI(Hep2064) by introduction of Sendai virus carrying the 4 Yamanaka factors [25]. The iPSC-O cells were cultured on mouse embryonic fibroblasts (MEFs) and proliferated using a medium for human ES cells [KNOCKOUT DMEM (Gibco,10829), 15% CTS KnockOut SR XenoFree Medium (Gibco,14150), 1% penicillin streptomycin (Gibco,11140), 1% Non Essential Amino Acid (Gibco,11360), 1% Sodium Pyruvate (Gibco,11360), 1% Gulta MAX (Gibco, 35050), 0.1% β-mercaptoetanol (Gibco,21985-023), and 10.0 ng/ml bFGF (INVITROGEN, PHG0024)].

### Preparation of feeder cells

Mouse embryonic fibroblasts (MEF) were prepared for use as nutritional support (feeder) cells. E12.5 ICR mouse fetuses (Japan CLEA) were taken out and the fetus head, limbs, tail, and internal organs were all removed, minced with a blade, and seeded in a culture dish in a medium (DMEM medium containing 10% FBS, 1% P / S) to allow cell growth. Using the X-ray irradiation apparatus (Hitachi, MBR-1520 R-3), 1/100 amount of 1 M HEPES Buffer Solution (INVITROGEN, 15630-106) was added to the obtained cells. Following irradiation with X rays (dose: 30 Gy), the obtained cells were frozen using a TC protector (DS Pharma Biomedical, TCP-001) and used as feeder cells.

### Hepatocytic differentiation and subsequent propagation

To generate embryoid bodies (EBs), ESCs and iPSCs (1 × 10^4^/well) were dissociated into single cells with accutase (Thermo Scientific, MA, USA) after exposure to the rock inhibitor (Y-27632: A11105-01, Wako, Japan), and cultivated in the 96-well plates in the EB medium [76% Knockout DMEM, 20% Knockout Serum Replacement (Life Technologies, CA, USA), 2 mM GlutaMAX-I, 0.1 mM NEAA, pen-strep, and 50 μg/mL l-ascorbic acid 2-phosphate (Sigma-Aldrich, St. Louis, MO, USA)] for 10 days. The EBs were transferred to the 24-well plates coated with collagen type I, and cultivated in the XF32 medium [85% Knockout DMEM, 15% Knockout Serum Replacement XF CTS (XF-KSR; Life Technologies), 2 mM GlutaMAX-I, 0.1 mM NEAA, Pen-Strep, 50 μg/mL L-ascorbic acid 2-phosphate (Sigma-Aldrich, St. Louis, MO, USA), 10 ng/mL heregulin-1β (recombinant human NRG-beta 1/HRG-β1 EGF domain; R&D Systems, Minneapolis, MN, USA), 200 ng/mL recombinant human IGF-1 (LONG R^3^-IGF-1; Sigma-Aldrich), and 20 ng/mL human bFGF (Life Technologies)] for 14 to 35 days. The differentiated cells were further cultivated and propagated in the ESTEM-HE medium (GlycoTechnica, Ltd., Japan) containing Wnt3a and R-spondin 1 at 37°C in a humidified atmosphere containing 95% air and 5% CO2 [26,27]. When the cultures reached subconfluence, the cells were harvested with Trypsin-EDTA Solution (cat#23315, IBL CO., Ltd, Gunma, Japan), and re-plated at a density of appropriately 5 × 10^5^ cells in a 100-mm dish. Medium changes were carried out three times a week thereafter.

### Selection with ammonia

Cells were treated with ammonia for 2 days to obtain ammonia-selected cells. The ammonia-selected cells were propagated for 7 days after exposure to ammonia in the modified F medium [28] until semi-confluence, and then passaged to 10-cm dishes after trypsinization with 0.25% trypsin/EDTA. In cases that number of cell death after ammonia treatment was high, the medium was changed immediately. In this study, ESC-cells and iPSC (iPSC-O) cells were exposed to ammonia.

### Hepatocytic maturation via spheroid formation

The ammonia-selected hepatocyte-like cells were detached with 0.25% trypsin/EDTA, transferred to 6-well plates at a density of 60,000 cells/well, and then cultivated for 7 or 10 days to form spheroid.

### Population doubling assay

Cells were harvested at sub-confluency and the total number of cells in each well was determined using the cell counter. Population doubling (PD) was used as the measure of the cell growth rate. PD was calculated from the formula PD=log_2_(A/B), where A is the number of harvested cells and B is the number of plated cells.

### Immunocytochemical analysis

Cells were fixed with 4% paraformaldehyde in PBS for 10 min at room temperature. After washing with PBS and treatment with 0.1% Triton X in PBS for 10 min, cells were pre-incubated with blocking buffer (5% goat serum in PBS) for 30 min at room temperature, and then exposed to primary antibodies [CYP3A4 (HL3, SANTACRUZ BIOTECHNOLOGY, sc-53850), α-Fetoprotein (R&D SYSTEMS, MAB1368), Human Albumin (CEDARLANE, CLFAG2140), Cytokeratin 7 (DAKO, M7018)] in blocking buffer overnight at 4°C. Following washing with 0.2% PBST, cells were incubated with secondary antibodies; either anti-rabbit or anti-mouse IgG conjugated with Alexa 488 or 546 (1:300) (INVITROGEN) in blocking buffer for 30 min at room temperature. Then, the cells were counterstained with DAPI and mounted.

### Quantitative RT-PCR

RNA was extracted from cells using the RNeasy Plus Mini Kit (QIAGEN: 74136). An aliquot of total RNA was reverse transcribed using an oligo (dT) primer (SuperScript TM □ First-Strand Synthesis System, INVITROGEN). For the thermal cycle reactions, the cDNA template was amplified (QuantStudio TM 12K Flex Real-Time PCR System) with gene-specific primer sets (Table 1) using the Platinum Quantitative PCR SuperMix-UDG with ROX (11743-100, INVITROGEN) under the following reaction conditions: 40 cycles of PCR (95°C for 15 s and 60°C for 1 min) after an initial denaturation (50°C for 2 min and 95°C for 2 min). Fluorescence was monitored during every PCR cycle at the annealing step. The authenticity and size of the PCR products were confirmed using a melting curve analysis (using software provided by Applied Biosystems) and gel analysis. mRNA levels were normalized using ubiquitin or GAPDH as a housekeeping gene.

**Table 1.**
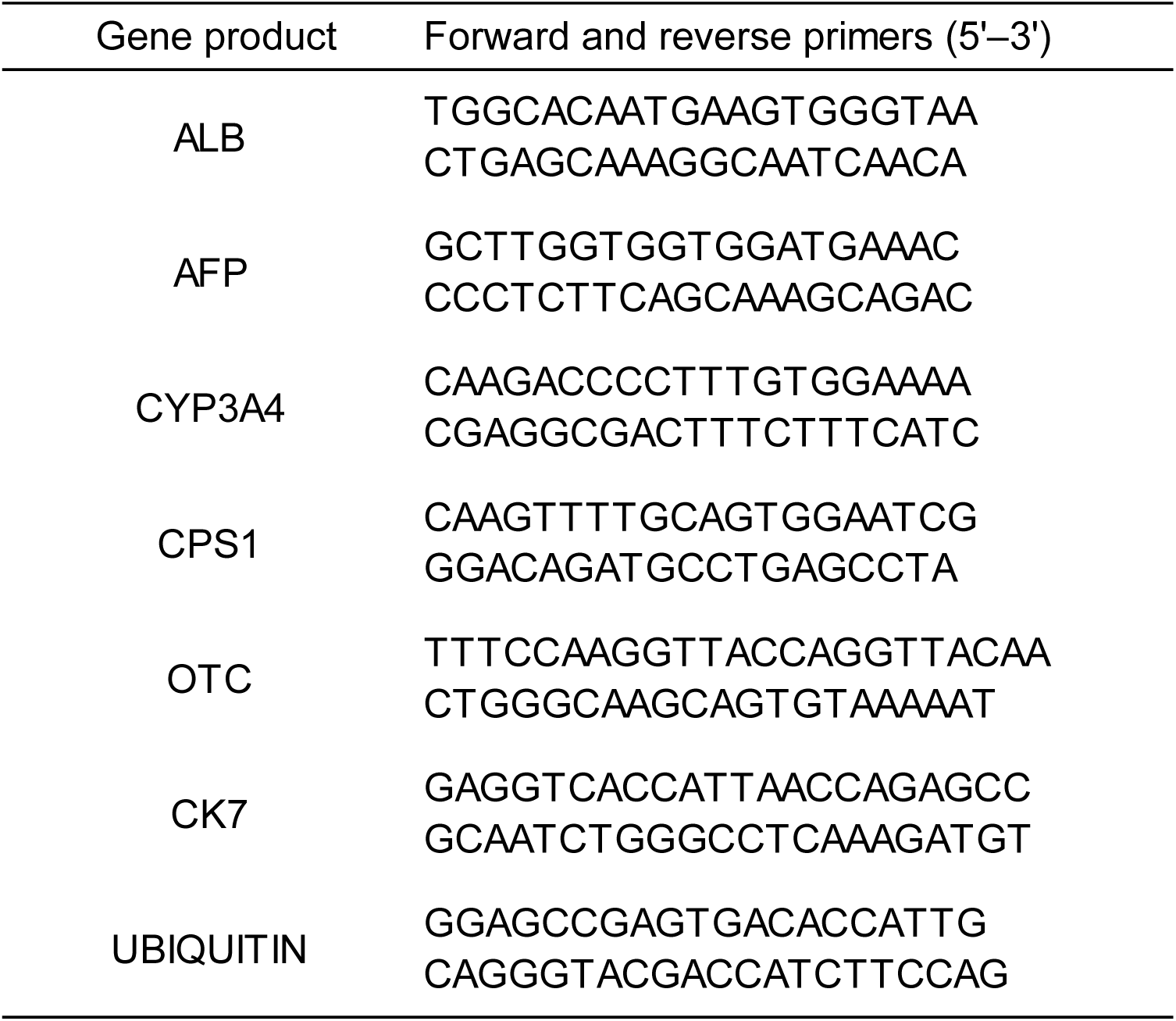
Primer pairs for RT-PCR

### Statistical analysis

Statistical analysis was performed using the unpaired two-tailed Student’s t test.

## Acknowledgements

We would like to express our sincere thanks to K. Miyado and H. Akutsu for fruitful discussion, to M. Ichinose for providing expert technical assistance, to C. Ketcham for English editing and proofreading, and to E. Suzuki and K. Saito for secretarial work.

